# A single light-responsive sizer can control multiple-fission cycles in *Chlamydomonas*

**DOI:** 10.1101/648436

**Authors:** Frank S. Heldt, John J. Tyson, Frederick R. Cross, Béla Novák

## Abstract

Proliferating cells need to coordinate cell division and growth to maintain size homeostasis. Any systematic deviation from a balance between growth and division results in progressive changes of cell size over subsequent generations. While most eukaryotic cells execute binary division after a mass doubling, the photosynthetic green alga *Chlamydomonas* can grow more than eight-fold during daytime before undergoing rapid cycles of DNA replication, mitosis and cell division at night, which produce up to 16 daughter cells. Here, we propose a mechanistic model for multiple fission and size control in *Chlamydomonas*. The model comprises a light-sensitive and size-dependent biochemical toggle switch that acts as a sizer and guards transitions into and exit from a phase of cell-division cycle oscillations. We show that this simple ‘sizer-oscillator’ arrangement reproduces the experimentally observed features of multiple-fission cycles and the response of *Chlamydomonas* cells to different light-dark regimes. Our model also makes testable predictions about the dynamical properties of the biochemical network that controls these features and about the network’s makeup. Collectively, these results provide a new perspective on the concept of a ‘commitment point’ during the growth of *Chlamydomonas* cells and hint at an intriguing continuity of cell-size control in different eukaryotic lineages.

**Graphical abstract:** 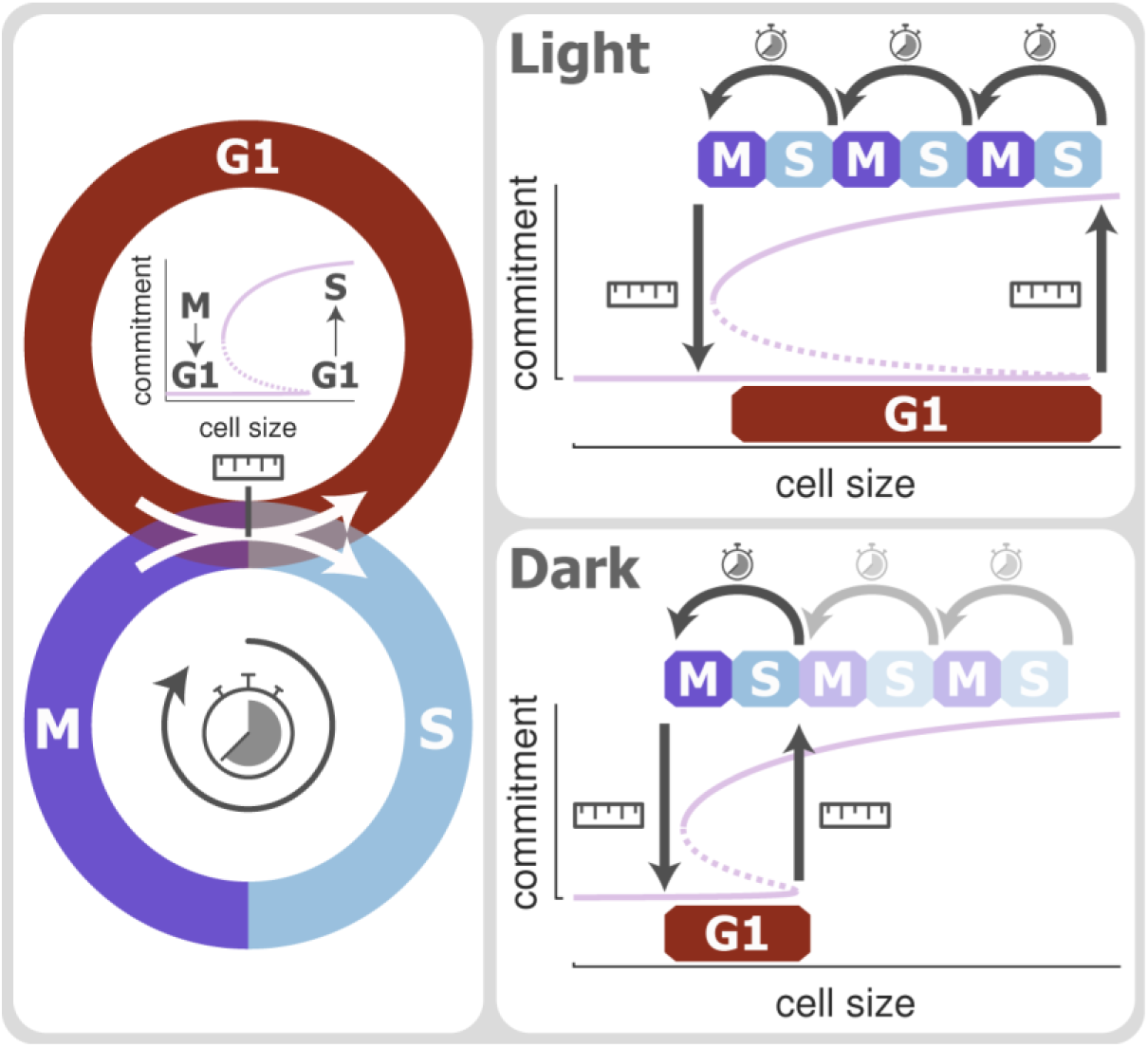

- G1-sizer and S/M-oscillator can give rise to multiple-fission cycles in *Chlamydomonas*
- Light-responsive bistable switch may guard transition between G1 and S/M-cycles
- Illumination increases S/M-entry threshold, causing multiple-fission cycles
- Dark shift lowers S/M-entry threshold, allowing small cells to commit to fewer divisions

## Introduction

Cell-cycle progression in Opisthokonts (fungi and animals) is governed by a common molecular mechanism based on cyclin-dependent kinases (CDKs) and their interaction partners: transcription factors, ubiquitin ligases, stoichiometric inhibitors, protein kinases and phosphatases [1–4]. Despite the universality of the general mechanism, there are subtle differences from one cell type to another in how the components interact and, consequently, in how cell-cycle progression is choreographed in time. For example, fission yeast cells have a short G1 phase and a long G2 phase, and they divide by binary fission. Budding yeast cells have a long G1 phase and a short G2 phase, and they divide asymmetrically. Acellular slime molds skip cell division altogether and produce multinucleate syncytia. Frog oocytes are arrested in G2 phase of meiosis I and grow to very large size; then, after fertilization, they undergo 12 rapid, synchronous S/M cycles to produce a hollow ball of 4096 small cells, from which the developing tadpole will arise.

In all these different cases, there are some common principles of eukaryotic cell-cycle progression. First of all, during mitotic cycles, S phase (DNA synthesis) and M phase (mitosis) alternate, so that mother and daughter cells maintain constant ploidy. There are rare exceptions to this rule, especially in some terminally differentiated tissues of multicellular organisms. Secondly, the average duration of the G1-S-G2-M division cycle (the cell cycle time) is identical to the mass doubling time (MDT) of the cell culture, so that cells maintain an ‘optimal’ DNA-to-mass ratio (within a factor of two) at all times. This rule also has its caveats: for example, mammalian cells seem to have a less strict ‘size control’ than yeast cells, and the average may need to be taken over a long period of growth and division (as in the case of oogenesis and early embryonic divisions). Thirdly, progression through the cell cycle is guarded by ‘cell cycle checkpoints’ that can block specific transitions (e.g. G1-to-S, G2-to-M and metaphase-to-anaphase) if problems are detected, such as DNA damage or misaligned chromosomes. Again there are caveats: not all checkpoints are operative in all cell types, and the early embryonic division cycles of frog eggs don’t seem to have any checkpoint controls at all [5]. It is full speed ahead for this very vulnerable stage of the frog life cycle.

A particularly interesting case is the unicellular green alga, *Chlamydomonas reinhardtii*, which grows during daylight to very large size (approximately 8-fold larger than its birth size), without undergoing DNA synthesis or cell division. Then, in the dark period of the daily rhythm, it executes a sequence of rapid cycles (*n* = 1, 2, 3 or 4, depending its growth during the light phase), each comprising S phase, M phase and cytokinesis, to produce 2^*n*^ progeny of a characteristic birth size. This pattern of growth and division is called ‘multiple fission’. Multiple fission most likely represents an evolutionary adaptation of green algae to maximize growth during the day, when photosynthesis is possible, while delaying the disruptive processes of DNA synthesis and cell division until night [6]. This mode of growth and division may be particularly advantageous for motile green algae like *Chlamydomonas* because they lose the ability of phototaxis when their flagella are resorbed prior to division, as their basal bodies are needed to coordinate mitosis and cytokinesis (an adaptation known as the ‘flagellation constraint’ [7]).

### Multiple-fission cycles in *Chlamydomonas*

The physiological properties of multiple-fission cycles in *Chlamydomonas* cells have been thoroughly studied by growing cells in different light:dark (LD) regimes (see Fig. 1A, based on [6,8,9]). In Fig. 1A, row (a), for instance, we illustrate the case of a 12:12 LD regime, which would be representative of algal cells under natural illumination. In the figure (which is a schematic representation of this common experimental culture condition), we assume that cells are growing with a 4 h MDT under nearly ideal conditions of illumination, temperature and nutrient availability. In this case, cells are growing in the light (a G1-type phase: growth without DNA synthesis or division) to a final size of approximately eight-times their initial birth size (Fig. 1B). Then, early in the dark phase, the ‘mother’ cell divides into eight daughter cells during three rapid S/M cycles (each with a period ≈ 30-40 min). For the remainder of the dark phase the daughter cells are in a G0-type phase (alive but not growing or replicating their DNA). After 24 h, at dawn, the cells resume growth (G0 to G1 transition), and the cycle repeats itself. In all other experiments in Fig. 1A, we assume that a cell culture is pre-synchronized by growth in 12:12 LD cycles.

**Figure 1.**
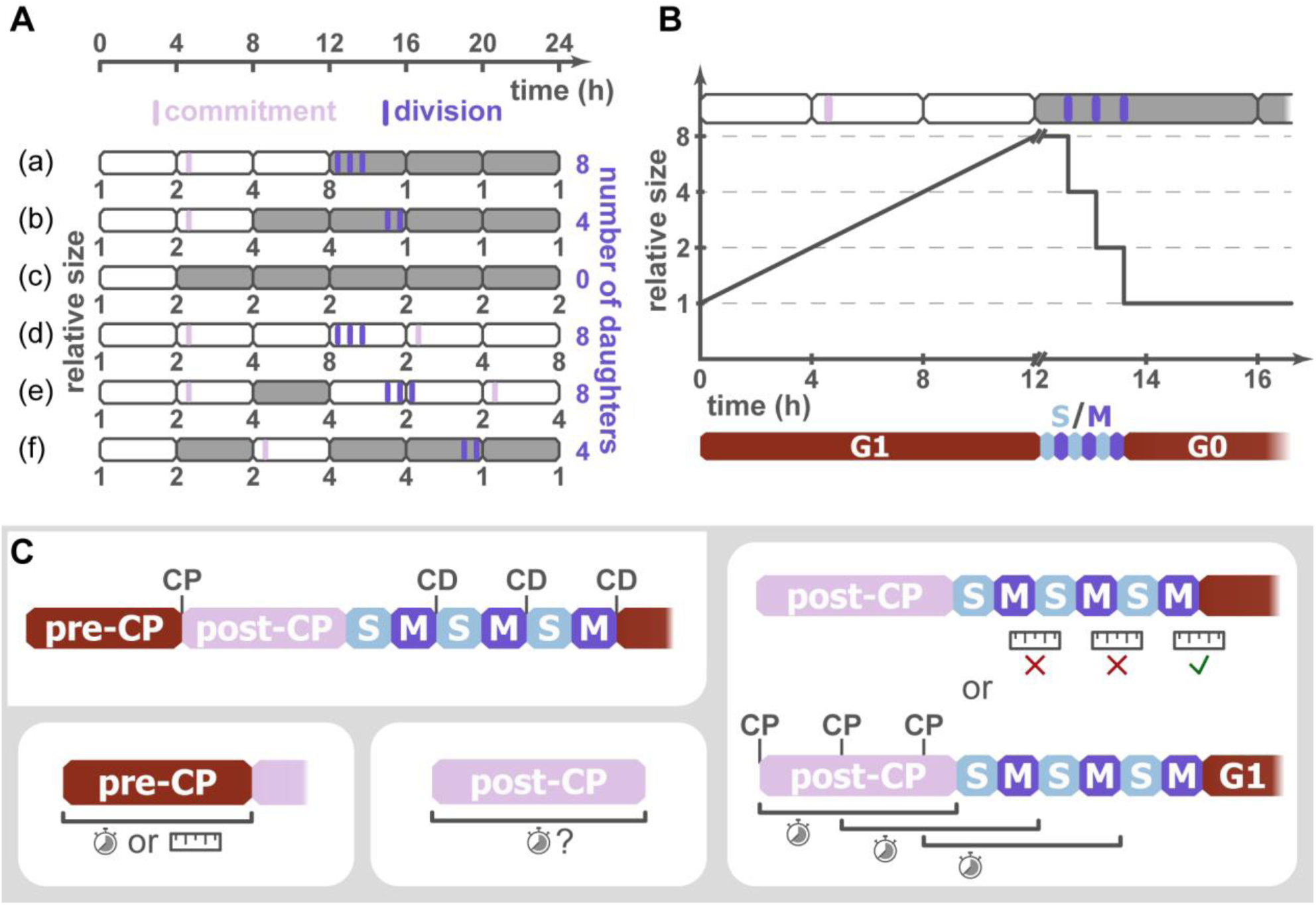
Multiple-fission cycles in *Chlamydomonas*. (**A**) Schematic of light-dependent growth and division. Four-hour intervals, equal to the assumed MDT, are represented by open (light periods) and closed (dark periods) rectangles. Cell size relative to birth size is shown below each row. Vertical stripes mark commitment and division. The final number of daughter cells is shown on the right. (**B**) Relative cell size during 12 h of light followed by a dark shift. Note the change in x-axis scaling after 12 h and the logarithmic y-axis. (**C**) Previous hypotheses for multiple fission. Cells commit to division in G1 phase at the commitment point (CP), but delay S/M cycles and cell division (CD). A timer or sizer might regulate commitment (bottom left), and timer control has been proposed for the post-CP period (bottom middle). To determine the number of divisions, cells might continue to divide until daughters fall below a threshold size (top right), or they might memorize the number of mass doublings they have undergone by acquiring serial commitments, each of which leads to division after a fixed delay (bottom right).

In Fig. 1A, row (b), the pre-synchronized cells are switched into constant darkness after 8 h of light to examine the mechanisms that provide specificity of division to darkness. At the beginning of the dark period, the mother cells are four-times their initial size, and, after a delay of ∼7 h, they undergo two S/M cycles to produce four daughter cells of size = 1. If this experiment is repeated with only a 4 h light period (row (c)), the mother cells are only twice the birth size, and they do not divide during the constant dark phase. These observations are commonly interpreted in terms of a ‘commitment point’ that a mother cell passes during its growth phase, after which completion of one or more cell-division cycles becomes independent of further growth [10–12]. Cells that are shifted into the dark (and stop growing) before the commitment point, will never divide. Cells that are shifted into the dark after this point divide at least once, after a long delay that depends on whether the cells are maintained in the light or shifted into the dark. For cells maintained in the light for a full 12 h (row (a)), the delay after commitment is ∼8 h. For cells growing in an 8 h light period, the delay is ∼10 h (row (b)). For cells grown in a 6 h light period (not shown), the delay is even longer.

What happens when cells are grown in constant light? As shown in row (d) of Fig. 1A, these cells pass the commitment point as in rows (a) and (b) and then divide in the light after ∼8 h, similarly to cells shifted to a dark phase at 12 h. In constant light, these cells undergo three rapid S/M cycles to produce eight daughter cells that are larger than the LD synchronized daughters, because (as we assume in this figure) they continue to grow in the light during the multiple-fission cycles. The eight daughter cells are in G1 phase (not G0) by definition (because they are growing), and they reach the next commitment point at *t* ≈ 17 h. The multiple-fission cycle time in constant light is 17 – 5 = 12 h, which is three MDTs and the multiple-fission *n* value is 3 divisions, to produce 2^3^ = 8 daughter cells, thereby maintaining balanced growth and division when averaged over a 12 h time window.

### Previous hypotheses to account for multiple fission in *Chlamydomonas*

The experimental observations in different lighting regimes sparked a number of proposals on how *Chlamydomonas* cells control their division cycle. Ultimately such hypotheses aimed to explain three characteristic features of multiple fission: (1) a cell commits to one or more divisions in G1, but (2) delays S/M cycles until long after the commitment point, and (3) a large mother cell divides into 2^n^ daughter cells, where *n* is ‘chosen’ such that the daughter cells have a characteristic birth size (Fig. 1C). Progression from cell birth to commitment was initially assumed to occur after a fixed time delay, controlled by a timer [12,13]. This mechanism was later revised to include a cryptic size dependence [6]. Subsequently, many experimental studies have suggested that cells need to reach a minimal size to attain commitment [8,11,14]. But there is no consensus on whether size is sufficient or a timer is involved as well [6]. Note that the commitment point is experimentally defined by shifting a culture into the dark [10–12]. Cells that divide sometime after the dark shift are deemed to have passed the commitment point, while those that do not are considered pre-commitment. To undergo multiple fission, cells must grow more than two-fold, and consequently they must suppress S/M cycles beyond the commitment point. Crucially, the prolonged period between commitment and S/M cycles is thought to be a distinct portion of G1, governed by different parameters [6,14]. This post-CP period was described by some as timer controlled [12,13], while others have argued for a variable post-CP period [9,15]. Finally, two hypotheses were proposed for how mother-cell size is linked to the number of divisions. The first assumes a mitotic sizer that allows cells to continue dividing after entering S/M cycles, until the daughter cells fall below a minimal size [6,11]. The second proposes a form of cellular memory that tracks the number of mass doublings a cell underwent by the attainment of multiple commitments [12,13]. In an extension of this mechanism [14], each successive commitment is assumed to license a subsequent division that is carried out after a fixed delay [14,16]. Note that this fixed delay runs contrary to some experimental observations. Specifically, each successive commitment would be attained after a mass doubling, suggesting that each successive S/M cycle would also follow after an interval equal to the MDT. Yet, S/M cycles are typically much shorter than and independent of the MDT [8]. Additionally, some studies indicate that brief periods of light support later divisions only after a considerable excess delay, i.e., the time delay is not fixed [11].

Little is known about how these complex controls could come about mechanistically. Intriguingly, though, genetic studies suggest involvement of a homolog of the animal retinoblastoma protein (Rb; MAT3 in *Chlamydomonas*), a transcriptional repressor, and its target transcription factor E2F/Dp, as well as a novel protein kinase CDKG1 that may regulate MAT3 [17–20]. In addition, *Chlamydomonas* cells show circadian rhythms in their behavior and physiology [21,22]. However whether a circadian clock controls the multiple-fission cycle remains controversial [6]. Several studies that varied the MDT time beyond traditional 24 h periods found no dependence of cell division on circadian rhythms [9,15,23,24], and, when kept under constant illumination, *Chlamydomonas* cells can adapt a variety of non-circadian division periods [8,9], arguing against control by a free-running clock.

In this study, we present a novel dynamical model of multiple-fission cycles in *Chlamydomonas*. While this model is based on known principles of eukaryotic cell-cycle control, it remains independent of the specific biochemical mechanisms operating in *Chlamydomonas*, which have yet to be fully uncovered experimentally. Our model suggests that temporal patterns of growth and division of *Chlamydomonas* cells can be reproduced by (1) a single sizer mechanism that guards the transitions from G1 phase into S/M cycles and from S/M cycles back to G1 phase, and (2) a cell-cycle oscillator that governs periodic progression through S/M cycles and cell division. This ‘sizer-oscillator’ arrangement provides new perspectives on the physiological concept of the ‘commitment point’ in *Chlamydomonas*. The model also suggests a conserved topology [4,25] of the biochemical networks that control cell size in Opisthokont and non-Opisthokont lineages.

## Results

### A sizer-controlled oscillator can explain multiple-fission cycles

Many eukaryotic cells delay progression through the cell-division cycle until they reach a critical size [26–29]. Based on this observation, we propose a simple model for multiple-fission cycles in *Chlamydomonas* (Fig. 2A). In our model, cells grow exponentially, provided that they are supplied with sufficient light and nutrients. A size-dependent biochemical regulatory network (a ‘sizer’) responds to increasing cell volume by activating a factor that induces competence for entry into division cycles. (Note that in the following sections, we use ‘cell volume’ and ‘cell size’ interchangeably.) By ‘division competence’ we mean transition to conditions permissive for the firing of a cell-cycle oscillator that spans S and M phases and directly controls genome replication and cell division. In this paper, we propose a specific biochemical implementation of the sizer and oscillator (Fig. S1A) based on the design principles of cell-cycle regulation that are known to be conserved in Opisthokonts [4]. Note however that the identities, interactions and indeed the very presence of many of these components are speculative in *Chlamydomonas*. The components and interactions in our model represent our best guess of what might be going on ‘under the hood’ of *Chlamydomonas* cells, but it is important to recognize that the general characteristics of the sizer-oscillator setup are independent of the specific biochemistry.

**Figure 2.**
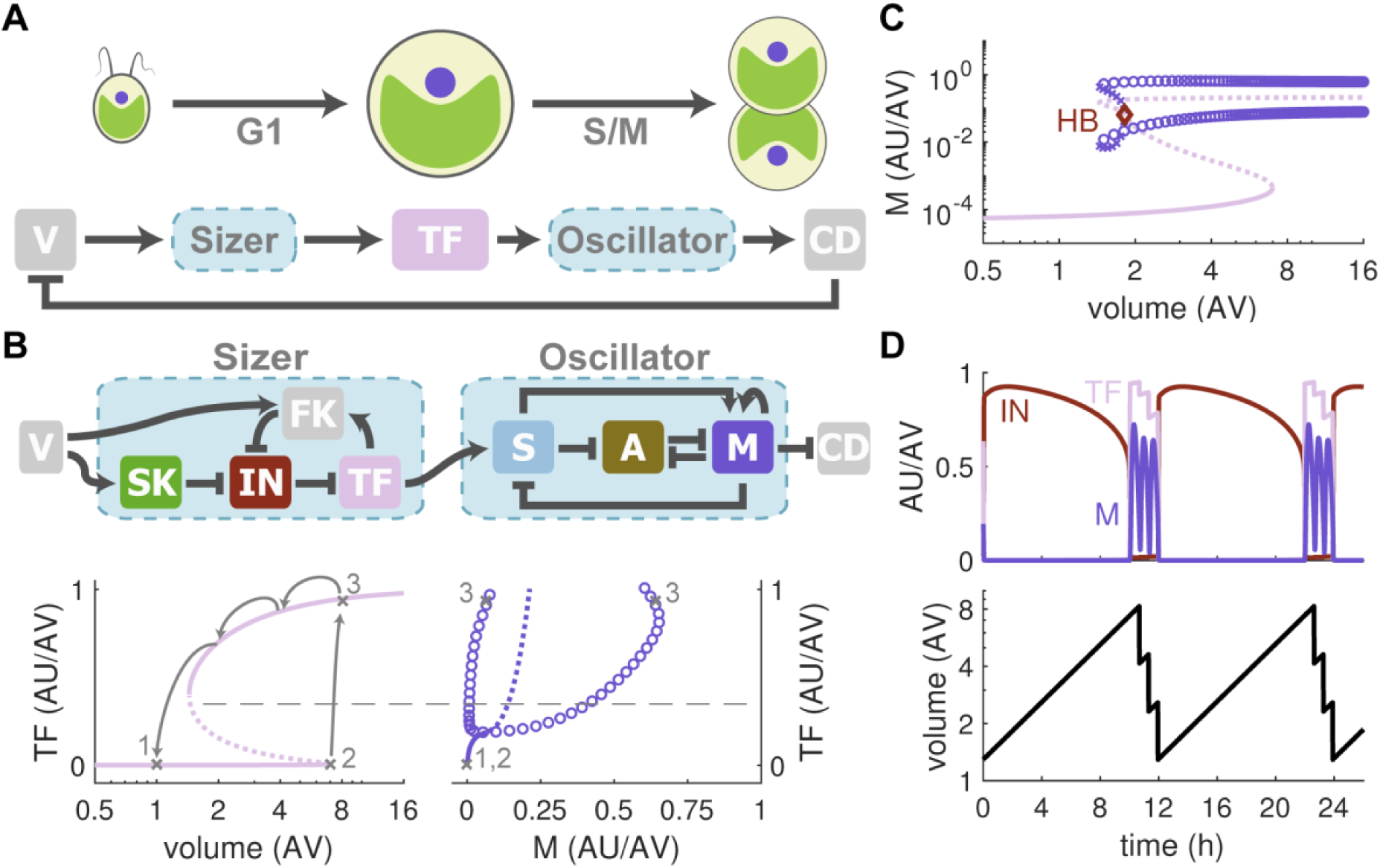
A sizer-oscillator arrangement can generate multiple-fission cycles. (**A**) Proposed mechanism for size control in *Chlamydomonas*. A biochemical network that guards cell-cycle entry acts as a ‘sizer’, converting information on cell volume (V) into the activity of a transcription factor (TF). Increased TF activity triggers an ‘oscillator’ that controls the events of S and M phase and initiates cell division (CD), which resets the system. (**B**) Molecular networks and signal-response curves of the sizer and oscillator. Solid and dashed lines indicate stable and unstable steady states, respectively. Open circles mark the amplitude of oscillations. Note that the oscillator diagram (bottom right) was turned 90° such that its input, TF, is shown on the y-axis to highlight equivalent TF values in the sizer output (bottom left). V, cell volume; SK, starter kinase; IN, inhibitor; TF, transcription factor; FK, feedback kinase; S, S-phase promoting factor; M, M-phase promoting factor; A, antagonist of M; CD, cell division. (**C**) Combined signal-response curve of the sizer-oscillator system. Red diamond marks the position of a Hopf bifurcation (HB). (**D**) Time courses of selected components (top) and cell volume (bottom) in a growing cell. AV, arbitrary unit of cell volume; AU/AV arbitrary unit of concentration.

In particular, we propose that the sizer (Fig. 2B, top left) is based on a transcription factor (TF) under the control of a stoichiometric inhibitor (IN). A starter kinase (SK) can inactivate IN, alleviating TF inhibition. Once TF becomes active, it promotes the production of a feedback kinase (FK) that further disengages IN from TF, creating a positive feedback loop. We assume that the levels of both SK and FK increase in direct proportion to cell volume. Differential scaling of protein concentrations with cell volume is commonly assumed to underlie size control [30,31]. While we do not specify the exact mechanism behind this proportionality here, elsewhere we have presented a mechanistic model for how such a scaling might arise [29]. Note that the sizer network is reminiscent of the Rb/E2F system in human cells and the Whi5/SBF system in budding yeast. In the well-established animal and budding yeast models, SBF and E2F activate transcription of multiple genes including cyclins, APC regulators and others, which together allow functioning of a cell-division-cycle oscillator. We propose that in *Chlamydomonas*, TF similarly promotes expression of such genes. Indeed, it is known that shortly before the multiple-fission cycle initiates in *Chlamydomonas*, there is massive transcriptional induction of essentially every gene known or suspected to be required for DNA replication, cell division, and operation of a cyclin-CDK oscillator [32–36]. Generically, the oscillator (Fig. 2B, top right) is based on interactions between an S-phase promoting factor (S) and an M-phase promoting factor (M). The synthesis of S is promoted by TF, and S inhibits an antagonist (A) of M. S also promotes the synthesis of M, which in turn downregulates S and A. Cell division occurs once the activity of M falls below a certain threshold, as the cell exits mitosis. The oscillator mirrors the interactions of cyclin-dependent kinases with the anaphase-promoting complex/cyclosome, and similar circuitry has been documented as acting in the *Chlamydomonas* division cycle [32]; but we emphasize, once again, that the dynamical behavior of the model (in both its sizer and oscillator modules) is independent of such speculative biochemical identifications.

First, we analyze the sizer and oscillator modules in isolation. With cell volume as its input, the sizer produces a bistable all-or-none response in TF activity (Fig. 2B, bottom left), which is typical for cell cycle entry [37,38]. Small cells show little or no TF activity, while TF activity is near maximal in large cells. TF activity in cells of intermediate size depends on their history. The oscillator’s input is TF activity. The oscillator sits at a stable steady state when TF activity is low (Fig. 2B, bottom right). But once TF reaches a critical level, oscillations in both S and M activities readily emerge. Combining these observations, we find that a small, newborn cell (point 1 in Fig. 2B) shows no TF activity and hence no oscillations in S and M. Strong inhibition of TF maintains this state as the cell grows to a volume almost eight-fold larger than its birth size (point 2). Shortly after that, further volume increase allows the starter kinase to overcome the inhibitor, which activates TF and triggers S and M oscillations (point 3). Because the cell now expresses FK, it can maintain TF activity even at lower cell volume. Hence, cell division, occurring at the end of each S/M-cycle, will not immediately result in TF inactivation. Only after the size of the daughter cells drops below a much smaller size threshold (after the third division in Fig. 2B) do the daughter cells exit the oscillatory regime and return to their initial G1 starting point 1.

Of crucial importance to our model, the bistable sizer switch has two distinct size thresholds because IN is inhibited by two different kinases. Small cells (in G1 phase) are reliant on the accumulation of SK to switch off IN and promote activation of TF. But, once TF is activated and the cell enters S/M cycles, FK is expressed and maintains IN in the ‘off’ state (and TF in the ‘on’ state) until cell size is reduced to a much smaller value (the threshold for TF inactivation). Combining these sizer and oscillator mechanisms allows newborn cells to remain in G1 up to a large cell volume (Fig. 2C). And then, after exceeding a critical size threshold (about eight-fold larger than their birth size), these cells transition rapidly into the oscillatory regime and undergo multiple cycles of S and M activities and cell divisions before returning to G1. Time course simulations of this model recapitulate the familiar pattern of multiple-fission cycles in *Chlamydomonas* observed under constant lighting (Fig. 2D and S1B). Specifically, cell volume increases during an extended G1 phase followed by three rapid division cycles, which produce eight daughter cells. The trajectory of these cells around the sizer and oscillator signal-response curves are shown in Fig. S1C. Collectively, these results demonstrate that coupling a cell-cycle oscillator to a cell size-dependent bistable switch with a large region of bistability (a stable G1 steady state coexisting with stable S/M oscillations; Fig. 2C) can create multiple-fission cycles characteristic of *Chlamydomonas* cells.

### A light-responsive sizer may control commitment

Since the growth of *Chlamydomonas* cells is highly dependent on illumination, we next consider the effects of variable light exposure in our model. Nutritional conditions are known to influence cell-cycle progression in other eukaryotes, where cells growing in rich medium delay cell-cycle transitions in order to grow to larger size [39,40]. These observations led us to assume that light, a crucial nutrient for *Chlamydomonas*, delays cell cycle entry. (This assumption is supported by experiments showing that light does indeed delay entry into multiple-fission cycles [41–43]. The effect is blue-light specific and delays the activation of S- and M-phase promoting factors; and the effect can be pharmacologically differentiated from the role of light in promoting photosynthesis). In our model, we assume that light promotes starter kinase degradation (Fig. 3A, top panel), as many nutritional signaling pathways converge on early cell-cycle kinases [44,45]. This light-responsive sizer shows no change in the all-or-none response of TF activity, with low activity in small cells and near-maximal activity in large cells (Fig. 3A, bottom panel). However, the critical size for entering S/M-cycles is significantly lower in the dark than in the light. More precisely, we assume that in the absence of light the starter kinase accumulates rapidly, allowing small cells to activate TF and leave G1 phase (Fig. S2A); whereas in the presence of light the starter kinase is degraded faster, and (consequently) cells must grow to a larger size to accumulate enough SK to activate TF. In the following discussions we refer to the critical size at which the G1-to-S/M transition occurs as the ‘TF activation threshold’. The ‘TF inactivation threshold’ refers to the opposite transition (S/M-to-G1), which occurs at critical sizes that are similar in both light and dark conditions.

**Figure 3.**
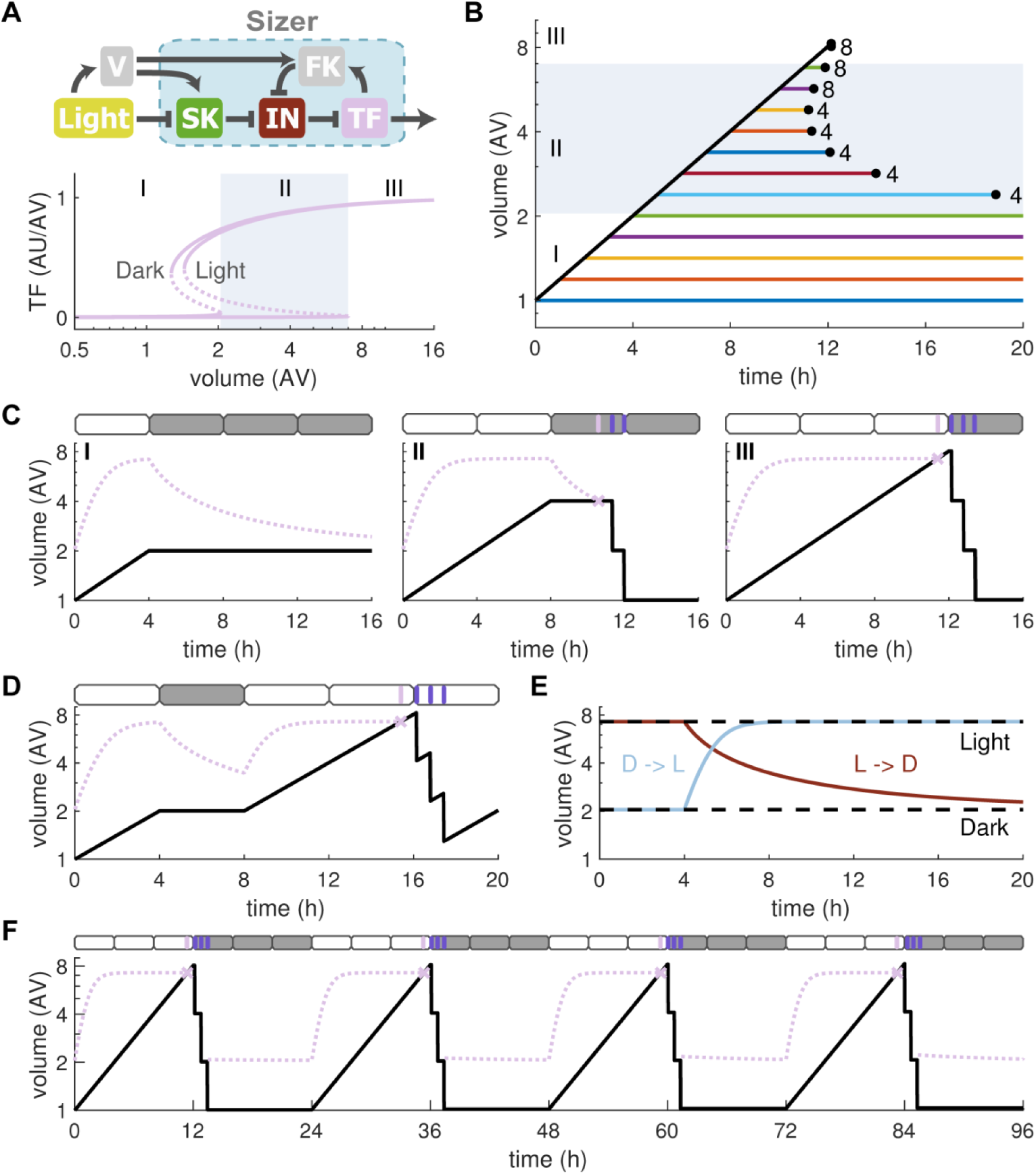
A light-dependent sizer can regulate commitment. (**A**) Molecular network and signal-response curves for a light-dependent sizer. Roman numerals correspond to classes of cells in characteristic size ranges. Shaded region marks Class II cells. (**B**) Cell size after varying periods of illumination. Black diagonal line corresponds to a cell in constant light. Colored horizontal lines show cells shifted into darkness after increasing periods of illumination (bottom to top: 0 to 12 h light period in 1 h steps). Black dots indicate time of first division and numbers indicate the final count of daughter cells. Cell classes from panel A are indicated. (**C**) Cell size over time for cells experiencing 4 h (I), 8 h (II) and 12 h (III) of illumination before a dark shift. Dashed lines indicate critical size for TF activation. (**D**) Size over time for a cell experiencing a 4:4:12 LDL illumination regime. Dashed line indicates critical size for TF activation. (**E**) TF activation threshold in light (upper dashed line) and dark (lower dashed line). Dynamics of threshold changes are shown for a dark-light (D→L) and a light-dark (L→D) shift after 4 h. (**F**) Cell size over multiple generations during 12:12 LD cycles. Dashed line indicates critical size for TF activation.

Based on these assumptions, three different classes of cells can be distinguished (represented in Fig. 3A by size regions I, II and III). Cells in Class I are below the TF activation thresholds in both light and dark. If subjected to a dark shift they stop growing and do not undergo S/M-cycles and cell division (Fig. 3B and C, region I). Class I cells are equivalent to ‘pre-commitment’ cells in the experimental literature [10–12]; our model suggests specifically that they are too small to activate TF even after prolonged darkness. Class III cells are large enough to pass the TF activation threshold even in the presence of light. They undergo multiple rounds of division with or without continued illumination (Fig. 3B and C, region III). Class II cells lie between these two extremes. They are not yet large enough to activate TF in the presence of light. However, when subjected to a dark shift, the TF activation threshold slowly decreases as the starter kinase re-accumulates (Fig. 3C, dashed lines). When the threshold falls below the size of a Class II cell, the cell activates TF, enters S/M-cycles and undergoes one or more cell divisions (Fig. 3B and C, region II). The number of daughters such a cell produces depends on its size (Fig. 3B). When the phase of multiple divisions reduces the size of daughter cells below the TF inactivation threshold, the daughters stop S/M cycling and enter G1 phase. By this mechanism, Class II cells can produce two, four or eight daughters after a dark shift (Fig. S2B). In addition, cells show a characteristic delay between the dark shift and the first division, which depends strongly on cell size at the time of the shift (Fig. S2B). A small cell, which is far below the TF activation threshold in the presence of light, experiences a long delay before the threshold has dropped enough to allow the cell to exit from G1 phase. The model also recapitulates the observation that, if growth is interrupted by a dark shift before a cell passes the ‘commitment point’ (i.e., grows larger than the TF activation threshold in the dark), it pauses in G1 but can resume growth and cell-cycle progression when illumination recommences (Fig. 3D). Together, these simulations are in excellent agreement with the experimental results summarized in Fig. 1A.

Since light affects the degradation of starter kinase in our model, changes in the TF activation threshold are governed by two characteristic time scales in response to changes in illumination. A light-dark (L→D) shift leads to a slow decrease in threshold size as SK re-accumulates slowly, while a dark-light (D→L) shift causes a rapid increase in threshold size as SK is rapidly degraded (Fig. 3E). The slow time scale causes the characteristic delay in cell division of Class II cells after a L→D shift. The more rapid change after a D→L shift ensures that, if illumination recommences, cells that have not yet entered S/M cycles grow to the full size seen under continuous light (see Fig. 3D). With the light-responsive sizer, our model also reproduces the characteristic growth pattern of *Chlamydomonas* cells in 12:12 LD cycles (Fig. 3F and S1D). Here, cells grow eight-fold in the light period before dividing three times into eight daughter cells of equal size in the dark. In an asynchronous population, such LD cycles lead to the typical synchronization of cell-division cycles seen experimentally (Fig. S2C). In all these different lighting regimes, daughter cells are born at a similar size (Fig. S2D), because the TF inactivation threshold (in our model) changes only slightly between light and dark conditions.

In simulations to this point, we have assumed that MDT = 4 h in the light and that the light periods come in 4 h increments. What happens if MDT is shorter or longer than 4 h? In Fig. S3 we investigate this question. For a shorter MDT = 3 h (faster growth), cells in constant light are predicted to undergo multiple-fission cycles with a period of 12 h that produce 16 daughter cells. In a 12:12 LD regime, this growth and division pattern is interrupted by 12 h of non-growth during the dark phase. For a longer MDT = 5 h (slower growth), cells in constant light undergo multiple-fission cycles of a period 15 h with three divisions per cycle to produce 8 daughter cells. In a 12:12 LD regime, these cells show a more complex pattern of growth and division: alternating between three divisions in one dark period and two divisions in the next. These predicted patterns of multiple fission cycles could be easily tested by altering light intensity (and thereby MDT) during the light periods.

In summary, a single light-dependent sizer can reproduce several important characteristics of the *Chlamydomonas* cell-division cycle, including growth of newborn cells to a large cell size in light, the commitment of smaller cells to fewer divisions after a dark shift, and the division into a variable number of daughter cells with constant size. Moreover, the model suggests that cells do not become committed to S/M cycles in mid G1 phase but rather at the end of it, when TF is activated. This ‘late’ commitment point is determined by cell size at the TF activation threshold, which is a function of light exposure in our model.

### Perturbations to sizer mechanism influence commitment and daughter size

To better understand how the sizer operates, we next analyze perturbations to its components. In general, the size distribution of a cell population depends on the position of four size thresholds (Fig. 4A). The critical sizes for TF inactivation in dark and light (a and b in Fig. 4A, respectively) regulate daughter cell size. No daughter cell in the respective lighting regime is larger than these sizes, since cells above the threshold would undergo another S/M cycle and division. Similarly, the smallest daughter cell is half the threshold size as it originates from a mother cell just above the critical size. The threshold size for TF activation in the dark (c in Fig. 4A) marks the size after which a cell commits to at least one S/M cycle when subjected to a dark shift. Hence, it sets the maximum cell size observed after a prolonged dark period. The critical size for TF activation in light (d in Fig. 4A) is related to the maximum size that cells achieve under any lighting regime. Any cell that grows larger than this size undergoes cell division irrespective of the lighting regime. Cells can slightly exceed this size if they continue to grow during the transition and the first S/M-cycle.

**Figure 4.**
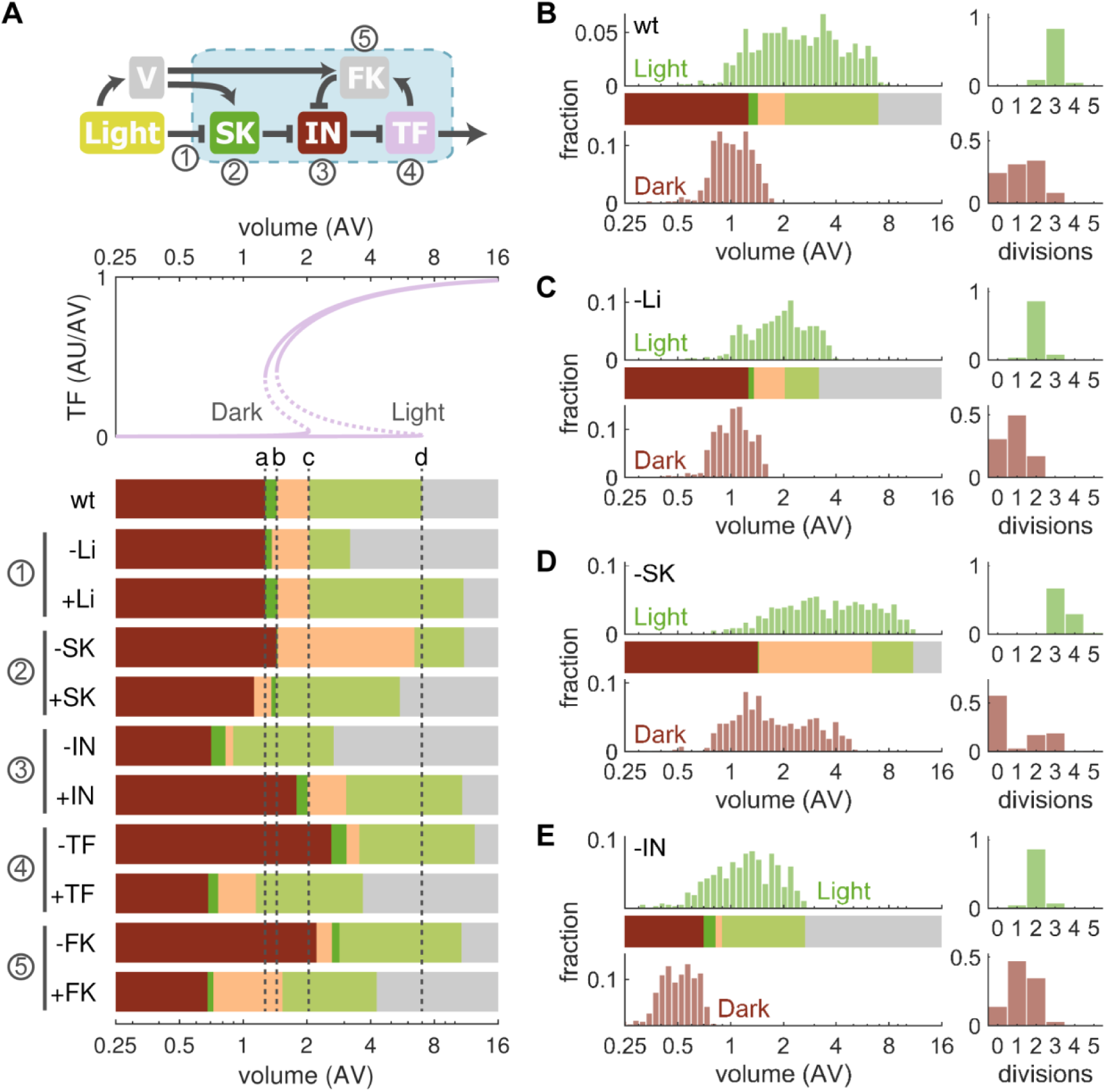
Effects of sizer perturbations. (**A**) How threshold sizes shift in response to perturbations of the sizer mechanism. The biochemical network (top) and signal-response curves (middle) for unperturbed (wild-type, wt) cells are shown to set a base-line for the effects of perturbations. Bars (bottom) mark the positions of the size thresholds for TF inactivation in dark and light (a and b, respectively) and TF activation in dark and light (c and d, respectively). Dashed vertical lines indicate the positions in wt cells. Numbers correspond to the mechanism or component in the schematic which is subjected to a reduction (-) or increase (+) of effect. (**B-E**) Distributions of cell size and division number in wild-type conditions (B), and for a reduction in light-dependent SK degradation (C), SK synthesis (D) and the inhibitor level (E). The size distribution of a cell population is shown after 14 generations in continuous light (Light) and at the end of a 24 h dark period that follows this continuous-light period (Dark). The number of cell divisions a mother cell undergoes in these regimes is shown on the right. Each panel comprises 500 individual cells from stochastic simulations. Bars from A are redrawn for comparison.

Under continuous light, an asynchronous wild-type (wt) population spans roughly an eight-fold size range, with cells dividing on average three times into eight daughters (Fig. 4B, top panels). Following a dark shift, up to three divisions are observed and cells revert back to smaller sizes and a narrower distribution; final cell sizes lie between the ‘c’ and half the ‘a’ threshold (Fig. 4B, ‘Dark’ panels). Perturbations in the light dependence of the sizer significantly change the ‘d’-threshold (Fig. 4A, -Li and +Li). For instance, a reduction in light-dependent SK degradation reduces the TF activation threshold during illumination such that cells reach smaller sizes and divide fewer times (Fig. 4C). However, the size distribution after a dark shift remains unaffected by this perturbation. For a change in SK synthesis, the model predicts a significant shift in the TF activation thresholds in both light and dark conditions (Fig. 4A, -SK and +SK). If SK production is compromised, size distributions change to larger sizes in both lighting regimes (Fig. 4D). Cells in continuous light tend to divide more often, while division numbers are reduced after a dark shift. Finally, perturbation of IN has a strong influence on all four thresholds (Fig. 4A, -IN and +IN). Reduced IN expression shifts cells to much smaller sizes and reduces division numbers (Fig. 4E). A decrease in the transcription factor (-TF) or feedback kinase (-FK) has the opposite effect on cell size (Fig. S4).

An interesting distinction between our model and the classical ‘commitment/delay’ hypotheses is that in the latter, there are three potentially fully independent mechanisms: (1) a sizer-dependent passage of ‘commitment’, (2) a timed blue-light-enhanced delay controlling the post-CP period, and (3) a second (mitotic) sizer allowing just the right number of divisions needed to return to a ‘target’ daughter size. In contrast, our model has two modules (sizer and oscillator) with shared components, leading to the prediction that most perturbations to the sizer affect several characteristic size thresholds simultaneously. Hence, mutations to the sizer control system can be expected to alter several features of multiple-fission cycles at the same time, and indeed this is observed [18,19].

## Discussion

Proliferating cells need to coordinate division and growth to maintain a near ‘optimal’ cell size (mass-to-DNA ratio) over multiple generations. Most cell types achieve this by undergoing binary division after a doubling of their mass. By contrast, the photosynthetic green alga, *Chlamydomonas*, grows more than two-fold in the daytime, putting off DNA synthesis and cell division until the night, when it divides multiple times to produce up to 16 daughter cells. In this work, we show that the characteristic features of this multiple-fission strategy for growth and division can be explained by a simple sizer that controls the growing cell’s entry into and exit from a phase of cell cycle oscillations. We provide evidence that the sizer is implemented by a bistable toggle switch with a large region of bistability, extending over an eight-fold range of cell volume. The ‘bistability’ is between a stable steady state corresponding to G1 phase of the cell cycle (growth but no DNA synthesis or mitosis) and stable limit cycle oscillations corresponding to multiple cycles of DNA synthesis, mitosis and cell division. That this bistable region extends over a large range of cell volumes ultimately enables the multiple-fission cycles so characteristic of *Chlamydomonas* cells.

Sizer mechanisms, whereby cells delay entry into the cell-division program (S and M phases) until they reach a critical size, control the START transition in budding yeast [26,46,47], as well as the G2/M transition in fission yeast cells [27,48,49] and slime mold plasmodia [28,50,51]. In previous publications, we showed that bistable toggle switches underlie these transitions in budding yeast [29] and fission yeast [52]. The mechanism controlling the G2/M transition in slime mold plasmodia is not so well characterized. It may be identical to the mechanism in fission yeast, based on competing effects of a kinase (Wee1) and a phosphatase (Cdc25). Alternatively, since the experimental studies referred to above suggest a mechanism based on the ‘titration of nuclear sites’, the G2/M transition in *Physarum* may be governed by a bistable switch analogous to the sizer in budding yeast [29].

The central insight proposed by our model for *Chlamydomonas* is that the same control principle can lead to multiple-fission cycles if the switch’s region of bistability spans a large range of cell volumes. Mechanistically, this requires strong inhibition of the G1-to-S/M transition, which constrains G1 cells to undergo more than one mass doubling, and a strong positive feedback that allows cells to maintain the S/M state once they have made the transition. In our model (Fig. 2B), the G1-to-S/M transition is induced by a transcription factor (TF). Strong inhibition of TF is provided by a stoichiometric inhibitor (IN) and the positive feedback is provided by a feedback kinase (FK) that is upregulated by TF and suppresses IN efficiently after cells have entered S/M cycles.

Guarding cell-cycle entry in this way provides several advantages to photosynthetic algae, such as *Chlamydomonas*. Firstly, the bistable switch allows for a phase of cell growth in the light and creates an all-or-none decision (with a clear size threshold) for S/M entry, beyond which the transition is irreversible. Hence, cells that pass this point commit to division and undergo at least one full cell cycle before returning to G1. Secondly, the switch sets a second, lower size threshold for S/M exit, which can regulate daughter cell size independently of commitment size. S/M cycles are maintained as long as cell size remains above this second threshold, leading to a constrained daughter size, regardless of the size increase experienced during the prolonged G1 growth phase. Thirdly, the number of cell divisions is variable, being determined by how many times a mother cell must divide to produce daughter cells that are smaller than the threshold size for S/M exit. Hence, a size-dependent bistable switch could account for the three main features of multiple-fission cycles, without invoking the more complicated mechanisms that have been proposed previously (see Fig. 1C).

Our proposal also suggests an intriguing continuity in eukaryotic cell-size control: both binary and multiple-fission cycles might rely on sizer-controlled oscillators that differ only in the sizer’s signal-response characteristics (Fig. 5). If the sizer shows a large bistable domain (Fig. 5, left panel), with the entry and exit thresholds for cell-cycle oscillations spanning more than a two-fold range of cell size (as we propose for *Chlamydomonas*), then multiple S/M cycles follow a long growth phase. However, binary division may suffice to return a cell to G1 phase if the bistable domain spans less than a two-fold range (Fig. 5, middle panel), or if the oscillator exerts a strong negative feedback on the sizer (Fig. 5, right panel). In the latter case (which we believe is characteristic of budding yeast cells [29]), entry into S/M cycles might switch off the sizer (e.g., by inhibiting the transcription factor), thereby forcing the sizer back into its G1 state after each division. In this case, the cell switches from G1 phase into S/M as a consequence of cell growth during G1 and from S/M phase back into G1 because the oscillator represses the sizer. That binary division and multiple fission might indeed use a similar control system is supported by the properties of early embryonic cell cycles, which seemingly transition between both modes. The developing frog egg, for instance, undergoes a long period of growth without division, before it starts a series of rapid, synchronous cell divisions without cell growth upon fertilization, resembling multiple S/M cycles in *Chlamydomonas*. Then, after the mid-blastula transition, subsequent cell divisions change to a more standard binary division pattern, as the developing embryo uses egg nutrient stores to support cell growth and division [5,30]. Formally, we propose a similarity between the extended growth of the frog egg before fertilization and rapid growth-independent divisions, and the multiple-fission cycle of *Chlamydomonas* following ∼8-fold cell growth during the long photosynthetically active G1.

**Figure 5.**
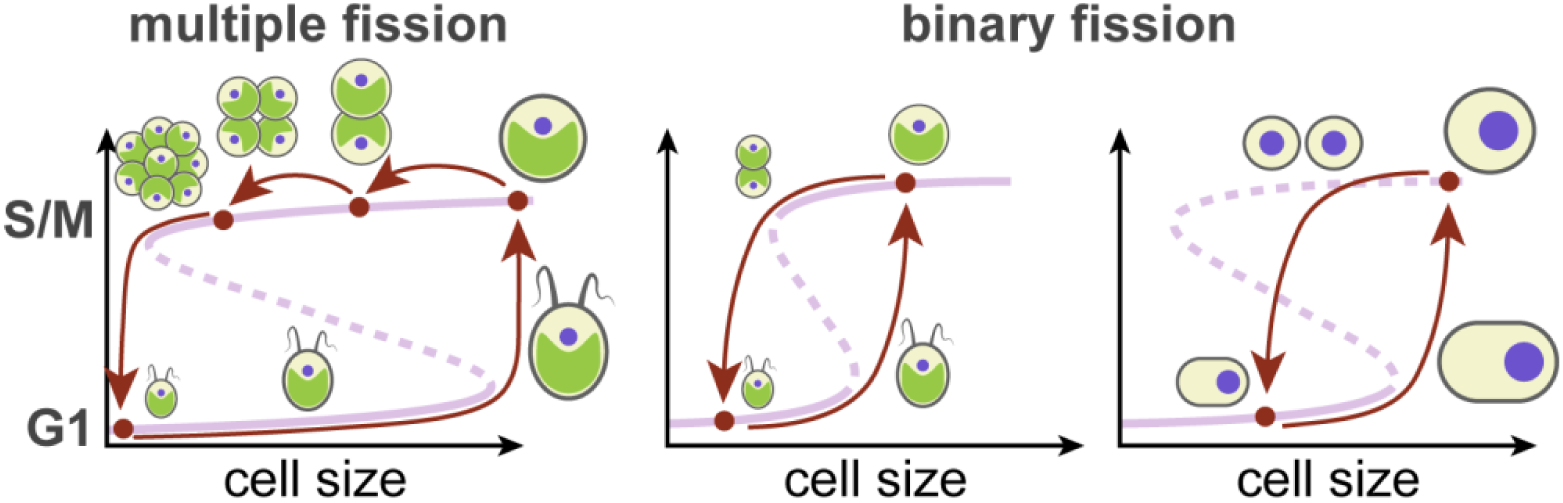
A sizer-controlled oscillator may account for both multiple and binary fission. (Left panel) A large bistable domain of the sizer, as proposed for *Chlamydomonas* in the light, leads to multiple-fission cycles. (Middle panel) A narrow bistability domain, as proposed for *Chlamydomonas* in the dark, may result in binary division. (Right panel) Strong negative feedback from the oscillator to the sizer, e.g. in budding yeast cells, forces the sizer into the ‘off-state’ after each division cycle, ensuring binary division.

Our model suggests that the varied responses of *Chlamydomonas* cells to LD shifts result from a direct influence of light on the sizer, which agrees with experimental observations where light exposure delays commitment and division [41–43]. Specifically, we propose that illumination affects the critical size for the G1-to-S/M transition, while hardly changing the threshold size of the reverse transition. This parallels the influence of nutrients on cell size in other organisms. Budding yeast cells, for instance, undergo START in rich medium at larger cell sizes [40], and fission yeast cells delay the G2-to-M transition under similar conditions [39]. Importantly, when nutrient supply suddenly worsens, cells respond with an accelerated transition, which allows smaller cells to complete the cycle [48]. We suggest that *Chlamydomonas* cells respond to a dark shift in a similar fashion, by reducing the critical size for S/M entry. While the exact molecular mechanism behind this change in *Chlamydomonas* cells remains to be elucidated, it may involve the loss of a cell-cycle inhibitor or accumulation of an activator following the dark shift.

A decreasing critical size for S/M entry in response to a dark shift could explain some of the more peculiar experimental findings in *Chlamydomonas*. Specifically, although cells are thought to pass a commitment point in mid G1, after which division becomes independent of growth, they experience (under constant illumination) a significant delay until they enter S/M cycles. The reason for this delay remains obscure and biochemical markers for commitment are lacking. Moreover, nearly every cell-cycle-related gene is upregulated just prior to or during S/M phase [32–36]. Our model attributes the delay to a requirement for cells to grow large enough to pass a ‘TF activation threshold’ near the end of G1 phase. Beyond this threshold, cells transit quickly and irreversibly from G1 phase to S/M cycles (i.e., it is at this point that they become biochemically committed to DNA synthesis and cell division). In light conditions, cells pass this threshold considerably later than the experimentally defined ‘commitment point’, and only shortly before the cell enters S phase. Hence, we propose that the previously defined pre- and post-CP periods are essentially indistinguishable from a cell-cycle perspective and that, although there are indeed many changes in gene expression during G1 phase [33,34,53], these changes are not causative agents for progression from G1 to S/M. Similarly, in our model, post-CP cells that are shifted into the dark become biochemically committed to division (activate TF) only after the S/M-entry threshold drops below their extant cell size. This occurs, we propose, at some time after the dark shift, with the delay depending on the cell’s size at the time of the dark shift: the smaller the cell, the longer the delay, exactly as observed experimentally [8]. The only measurable, cell cycle-related difference between pre-CP and post-CP cells immediately after a dark shift might be the loss of an inhibitor or accumulation of an activator of the cell cycle in post-CP cells (e.g., the accumulation of SK in our model).

Considering these findings, we propose a new interpretation of the ‘commitment point’ in the *Chlamydomonas* cell cycle. Rather than being a static control point that cells pass early in G1 and that is followed by a long and biochemically distinct post-CP phase [6], we propose that illumination simply delays S-phase entry by increasing the critical size for G1 exit, leaving much of G1 as a homogeneous growth phase with no discernable biochemical ‘commitments’ to DNA synthesis, mitosis and cell division. After a dark shift, cells remain in a similar (biochemically uncommitted) state until the decreasing size threshold drops sufficiently, at which point they commit to division.

In this study of multiple-fission cycles in *Chlamydomonas* cells, we present a simple mechanistic model based on ‘generic’ protein interactions (kinases, inhibitors, transcription factors, S- and M-phase promoting factors) without tying the model down to any specific — and necessarily speculative — identification of these proteins in *Chlamydomonas*. Nonetheless, our generic model bears many suggestive analogies to the consensus mechanism for cell-cycle control in Opisthokonts [1–4]. Recognizing that the model’s main conclusions are independent of its specific biochemical implementation and that assigning *Chlamydomonas* proteins to our generic regulators is a major challenge, we nonetheless venture to speculate about the biochemical makeup of the sizer and oscillator, based on some pioneering studies of cell-size mutants in *Chlamydomonas* [17–20]. Central to our model is the interaction of a stoichiometric inhibitor (IN) with a transcription factor (TF). We suggest that these functions may be provided by the products of the MAT3 (an Rb-homolog) and E2F1 genes, respectively. In support of this suggestion, a reduction in IN levels in our model causes a severe decrease in cell size, recapitulating the much-reduced daughter cell size in MAT3-mutant cells [18,20]. This effect resembles the cell-size reduction observed upon the deletion of Whi5, a stoichiometric inhibitor of the SBF transcription factor in budding yeast [54,55]. Conversely, loss of function of E2F1 and DP1 (an E2F-binding partner) leads to large cell-size phenotypes in *Chlamydomonas* [19], an observation reproduced by our model when simulating decreased TF levels. Although *mat3* and *e2f1* mutants show almost no changes in the overall gene transcription program during S and M phase [19], this observation does not preclude roles for MAT3 and E2F as specific transcriptional regulators (IN and TF) in our model; or, it may be that E2F1 promotes cell cycle progression in a transcription-independent manner [19,20] but still by providing a permissive environment for operation of the cell-division-cycle oscillator, as in our model. Considering the role of D-type cyclins as ‘starter kinases’ in other organisms, SK activity in our model could be provided by any of the four D-type cyclins in *Chlamydomonas* [6] and their CDK interaction partner(s). A recent study identified a cyclin-dependent kinase, encoded by CDKG1, that is expressed during S/M cycles and phosphorylates Mat3 [17]. This protein fits the description of the feedback kinase (FK) in our model. Indeed, a knockdown of CDKG1 leads to large daughter cells, similar to what we observe in simulations with reduced FK levels. Based on their roles for cell-cycle initiation, we propose that CDKA and its corresponding cyclin CYCA1 correspond to the S-phase promoting factor, while CDKB and CYCB1 are candidates for the M-phase promoting factor, considering they are essential for mitosis [6,32,35]. The antagonist in our model most likely corresponds to the *Chlamydomonas* variant of the APC/C, since experiments have established its antagonism to cyclin-CDKs [32].

Of note, cell-size mutants frequently show changes in both commitment size and division number in experiments [18,19]. This observation would be surprising if these two features were controlled by entirely distinct pathways. But it is in fact expected in our model, where commitment and division number are consequences of a single size-dependent switch. In the future, the increasing availability of *Chlamydomonas* mutants deficient in major cell-cycle regulators will help to refine the picture of multiple-fission control [35,53].

## Conclusion

Our results demonstrate that a single size-dependent bistable switch that guards the transition between a growth phase and the cell-division cycle can reproduce many of the characteristic features of multiple-fission cycles in *Chlamydomonas*. This sizer-controlled oscillator bears striking similarities to the size checkpoints found in other eukaryotes, suggesting a universal control paradigm for cell size. It also hints at a surprising malleability of the eukaryotic cell cycle, where modest changes to the control mechanism can generate diverse behaviors ranging from binary division to multiple-fission and embryonic cell cycles.

## Acknowledgments

FSH and BN are funded by the Biotechnology and Biological Sciences Research Council (BBSRC) strategic LoLa grant BB/M00354X/1. JJT acknowledges financial support from the US National Institutes of Health (5R01-GM078989) administered through Colorado State University (PI: Jean Peccoud). FRC is supported by the National Institutes of Health Grant 2R01-GM078153.

## Author contributions

Conceptualization: FSH, JJT, FRC and BN; Software: FSH and BN; Formal Analysis: FSH; Writing – Original Draft: FSH and JJT; Writing – Reviewing & Editing: FSH, JJT, FRC and BN; Visualization: FSH; Supervision: BN; Funding Acquisition: JJT, FRC and BN.

## Declaration of interests

The authors declare no competing interests.

## Model and methods

### Mathematical model

We model cell growth and multiple fission in *Chlamydomonas reinhardtii* using a set of ordinary differential equations. These equations describe the time evolution of cell volume and cell cycle regulators (see Fig. 2B and S1A). Growth of cell volume (*V*) is assumed to be exponential and light dependent [8,9]

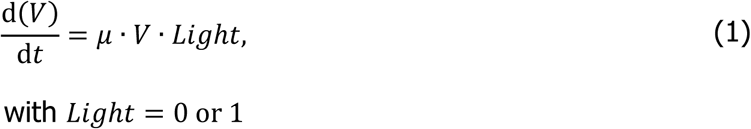

where *μ* is the maximum specific growth rate and *Light* is a binary variable for the influence of light on cell growth. For simplicity, we assume that growth occurs with maximum rate under illumination and ceases in the dark. More complex growth dynamics, which include the influence of light intensity [8,9], nutrients or temperature [12,15], may be of interest in future studies.

Rate equations for the molecular components in Fig. 2B are derived from the more detailed ‘biochemical reaction diagram’ in Fig. S1A. The concentration of the starter kinase (*SK*) changes according to

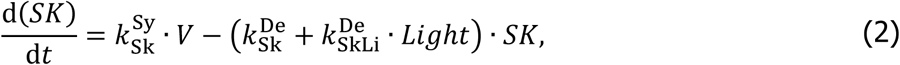

where synthesis occurs with rate 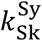 and increases with cell volume. While we here choose to implement the volume dependence of starter kinase synthesis (and feedback kinase synthesis, below) by simple volume multiplication, we have presented a more detailed, mechanistic model of how protein levels scale with volume in an earlier publication [38]. Starter kinase degradation occurs at a constitutive rate 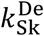 and a light-dependent rate 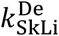, which reduces SK levels under illumination. We have chosen to degrade the starter kinase in the light because light represents a critical nutrient for *Chlamydomonas* and nutrient signaling typically impacts early stages of the cell cycle [44,45].

To describe the dynamics of the transcription factor and its inhibitor, we assume, first of all, that they are both present at a fixed total concentration (*TF*_t_ and *IN*_t_, respectively) in the cell

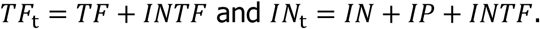

Free transcription factor (*TF*) and inhibitor (*IN*) bind tightly to form an inactive complex (*INTF*), as described by the first two terms in the following equations:

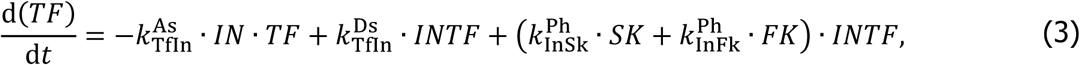

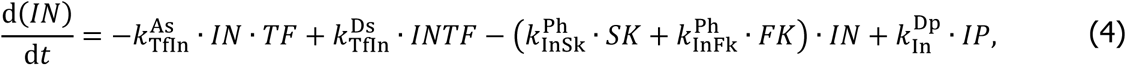

where the association and dissociation rate constants are 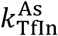 and 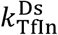, respectively. The later terms in Eqs. (3) and (4) describe inhibitor phosphorylation by the starter kinase with rate 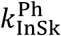 and by the feedback kinase (*FK*) with rate constant 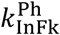. We assume that phosphorylation of the inhibitor disrupts its binding to TF. The phosphorylated inhibitor (*IP*) is dephosphorylated by a first-order reaction with rate constant 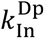. We assume that the feedback kinase is synthesized and degraded rapidly, such that it is always present at steady-state concentration in the cell. Synthesis of the feedback kinase is proportional to cell volume and promoted by the transcription factor. Hence, by absorbing the synthesis and degradation rates into the rate constant 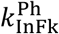, we can obtain the level of feedback kinase as

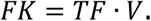

In our model, the S-phase promoting factor (*S*), M-phase promoting factor (*M*), and the antagonist (*A*) form a limit cycle oscillator governed by the equations

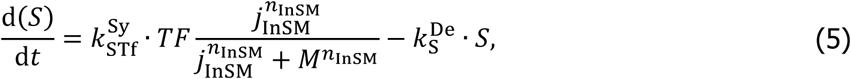

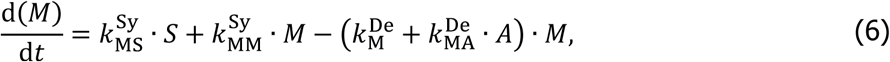

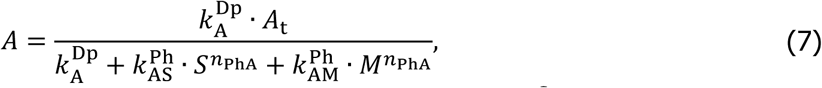

where the rate of synthesis of *S* is determined by a rate constant 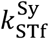, by the activity of the transcription factor and by the probability that the expression of the *S* is inhibited by *M*. The latter probability is given by a Hill function with half-saturation constant *j*_InSM_ and Hill exponent *n*_InSM_. Constitutive degradation of *S* occurs with rate constant 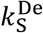. The rate of synthesis of *M* depends on *S* with rate constant 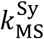 and in an autocatalytic manner on *M* itself with rate constant 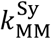. Degradation of *M* is comprised of a constitutive term with rate constant 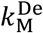 and an antagonist-dependent term with rate constant 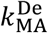. In Eq. (7) we are assuming that the total concentration of the antagonist (*A*_t_) is constant and that its active form (*A*) can be inactivated by both *S* and *M* through phosphorylation. The active and inactive forms of *A* are assumed to be in pseudo-steady state, with dephosphorylation (activation) occurring with rate 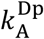 and phosphorylation (inactivation) by *S* and *M* occurring with rates 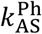 and 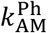, respectively. Phosphorylation depends on *S* and *M* in an ultrasensitive fashion with exponent *n*_PhA_.

Finally, we assumed that cell division ensues once *M* drops below a threshold concentration (*CdTh*), which is required to maintain the mitotic state, i.e.,

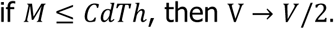

This cell division event produces two equally-sized daughter cells, one of which the model follows subsequently.

### Computation

The model was built within the Systems Biology Toolbox 2 [56] for MatLab (version R9.5.0 R2018b) and simulated using the CVODE routine [57]. Bifurcation diagrams were calculated within the freely available software XPPAUT [58]. The model will be made available in different formats after publication at www.cellcycle.org.uk/publication. Model files have also been deposited in BioModels [59] and assigned the identifier MODEL1904020001. Model parameters and initial conditions are listed in Tables S1 and S2, respectively, and Table S3 shows the parameter changes used to obtain the sizer perturbations in Fig. 4 and S4. In order to obtain size distributions and division numbers in Fig. 4B-E and S4, we simulated a stochastic version of the model using a custom-made MatLab code implementing Gillespie’s stochastic simulation algorithm (reviewed in [60]), according to the sorting direct method [61]. To this end, rate expressions of the deterministic model were converted into propensity functions assuming that 1 AU corresponds to 50 proteins and volume growth occurs according to Eq. 1 in increments of 0.01 AV.

