## Supplemental Information for "A single light-responsive sizer can control multiple-fission cycles in *Chlamydomonas*"

#### Contents

Supplemental figures ........................................................................................... 2
Supplemental tables............................................................................................. 6

### Supplemental figures

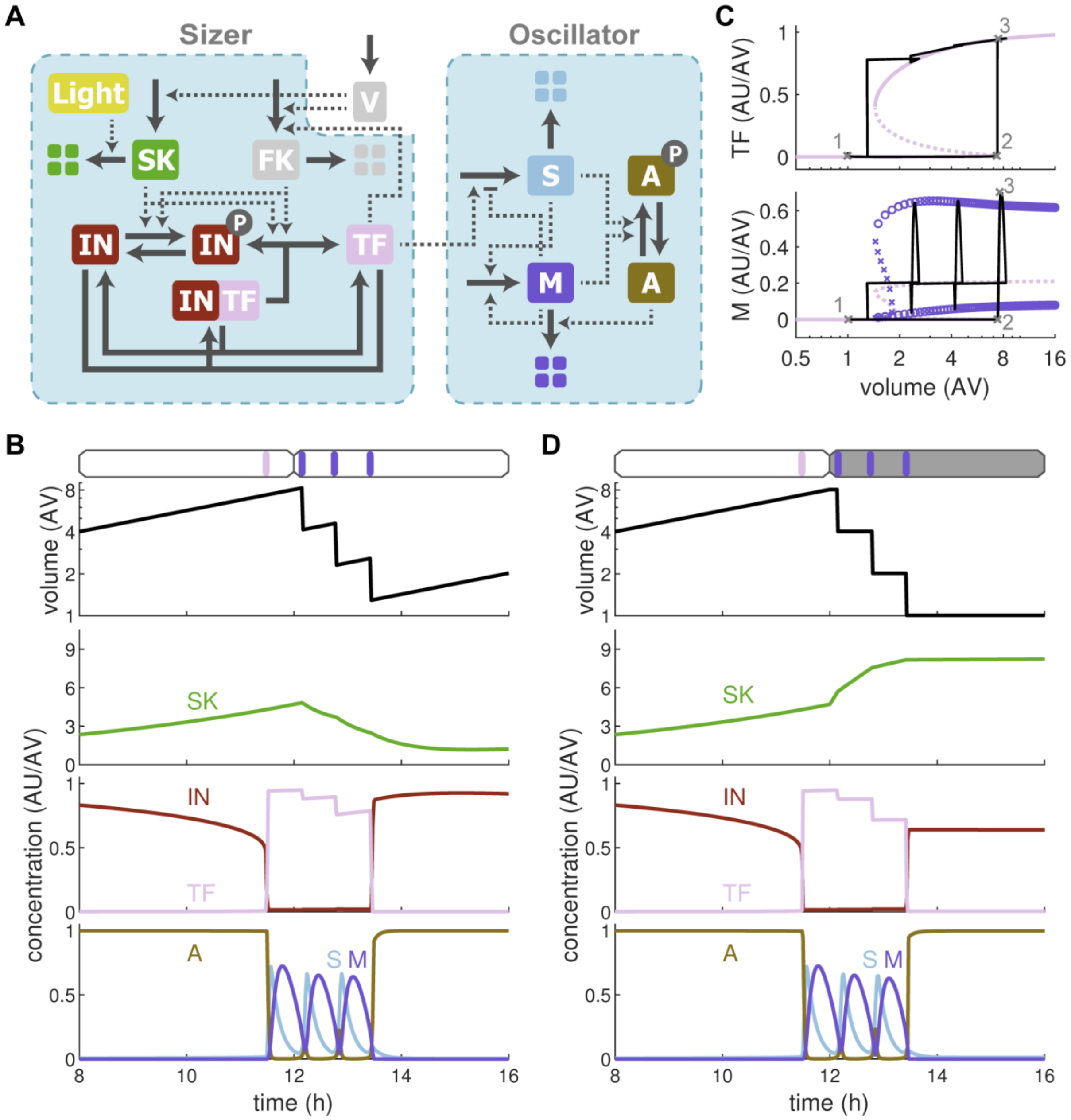


**Figure S1 | Related to Fig. 2.** (**A**) Biochemical reaction network of the multiple-fission model. Solid arrows represent synthesis, degradation, complex formation and dissociation, and phosphorylation and dephosphorylation reactions. Dashed lines with arrowheads and bars represent activation and inactivation, respectively. (**B, D**) Cell volume (top) and cell-cycle regulators (bottom) in constant light (B) and for a dark shift after 12 h (D). The time window between 8 and 16 h after the start of illumination is shown. (**C**) The trajectory of cells from Fig. 2D is shown as the black solid line, overlaid on the sizer diagram from Fig. 2B, lower left, the combined signal response curve from Fig. 2C.

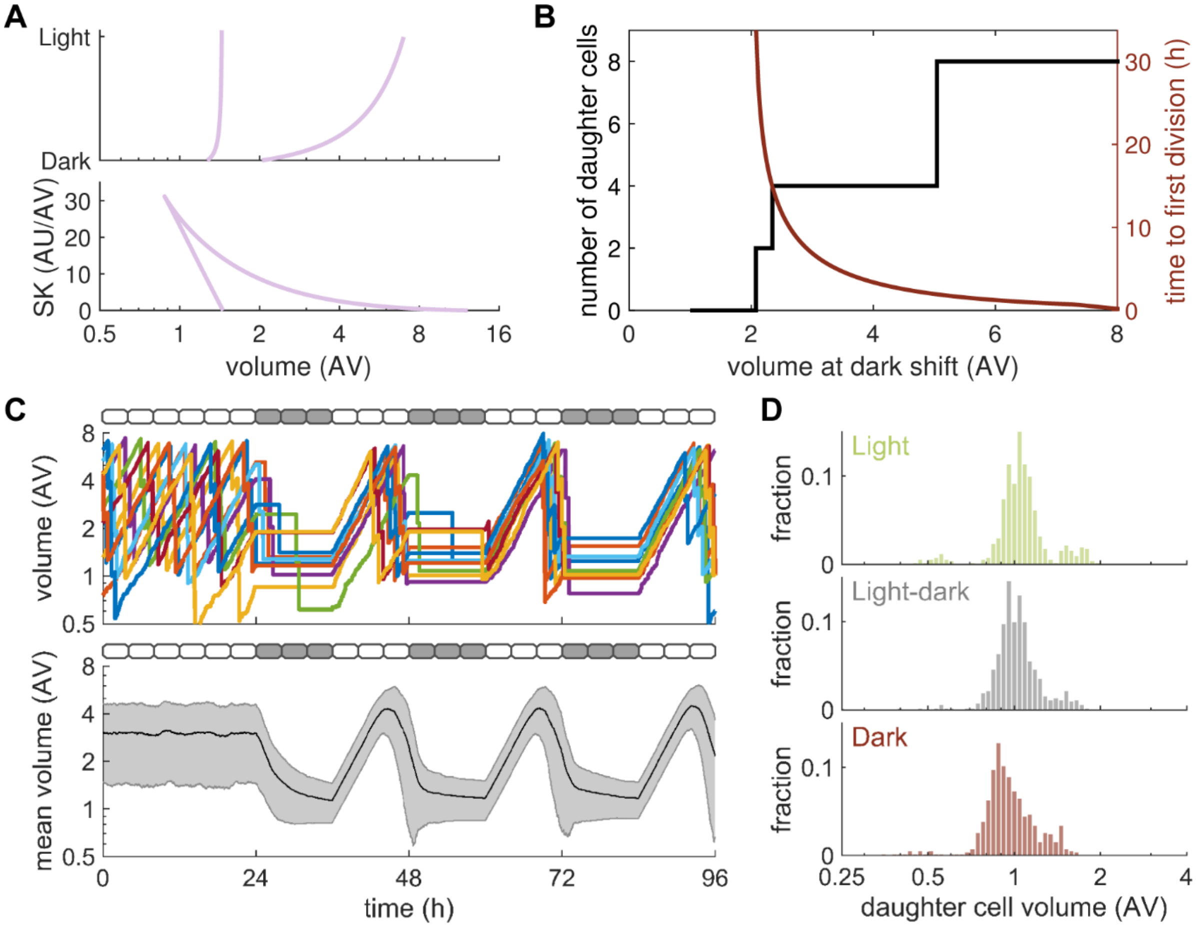


**Figure S2 | Related to Fig. 3.** (**A**) Dependence of the cell-volume thresholds for TF activation/inactivation (right branch and left branch, respectively) on light intensity (top) and on SK activity (bottom). (**B**) Number of daughter cells and time from dark shift to first division for cells of varying size at the time of the dark shift. (**C**) Stochastic simulations of 500 asynchronous cells experiencing 24 h of light followed by 12:12 LD cycles. Cell volume of five exemplary cells (top) and the mean cell volume and standard deviation (bottom) are shown. (**D**) Daughter cell size in 500 stochastic simulations under constant illumination (Light), during 12:12 LD cycles (Light-dark) and after prolonged darkness (Dark).

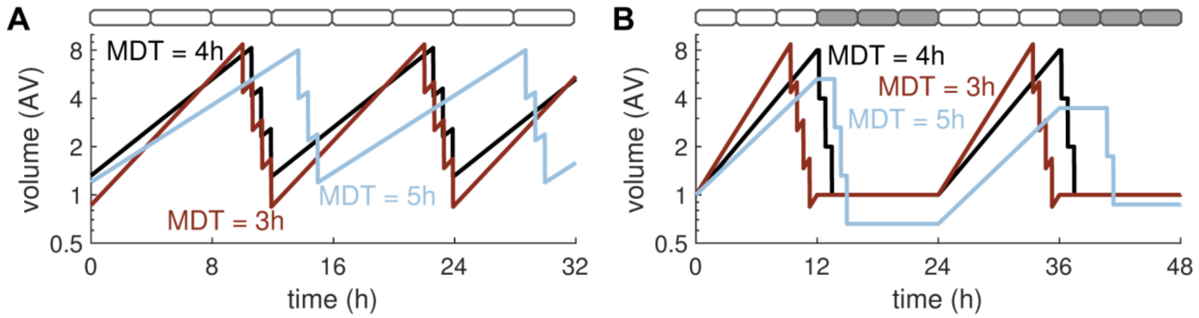


**Figure S3 | Related to Fig. 2 and 3.** Influence of MDT on multiple-fission cycles. Cell volume over time under continuous illumination (**A**) and during 12:12 LD cycles (**B**) for mass doubling times of 3 h, 4 h and 5 h. (**A**) In constant light, cells grown with MDT = 4 h undergo multiple-fission cycles with a period of 12 h and three divisions in each cycle. At MDT = 3 h, the mother cell still commences multiple-fission cycles around a volume of 8 AV, but it subsequently undergoes four divisions to produce 16 cells during a total cycle time of 12 h. For MDT = 5 h, three divisions are observed during a cycle time of 15 h. (**B**) In a 12:12 LD regime, cells grown and MDT = 3 h and 4 h show similar cycles than in panel A. But these cycles are interrupted by a 12 h period of no-growth in the dark. Cells grown at MDT = 5 h alternate between two and three divisions in the dark because 24 h of growth are nearly five mass doublings. Cells at MDT= 5 h could still be synchronized effectively over a series of 12:12 LD cycles, but 15:9 LD cycles would synchronize them more efficiently (simulations not shown).

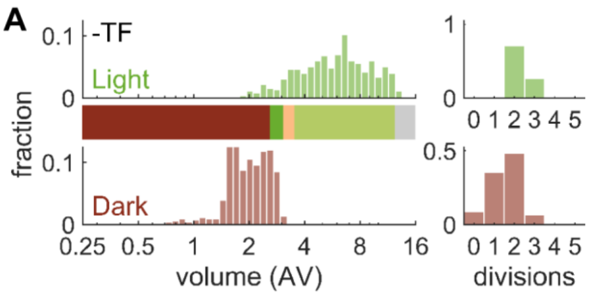


**Figure S4 | Related to Fig. 4.** Cell size distributions and division numbers for (**A**) a reduction in TF and (**B**) a reduction in FK. The size distribution of a cell population is shown after 14 generations in continuous light (light) and at the end of a 24 h dark period that follows this continuous-light period (dark). The number of cell divisions a mother cell undergoes in these regimes is shown on the right. Each panel comprises 500 individual cells from stochastic simulations. Bars from Fig. 4A are redrawn for comparison.

### Supplemental tables

**Table S1. Parameters of the multiple fission model.**

| Parameter | Description | Value | Unit[a] |
|---|---|---|---|
| $\mu$ | specific growth rate | 0.0029 | 1/min |
| $A_{\mathrm{t}}$ | total antagonist concentration | 1 | AU/AV |
| $CdTh$ | cell division threshold for $M$ | 0.2 | AU/AV |
| $IN_{\mathrm{t}}$ | total inhibitor concentration | 2 | AU/AV |
| $j_{\mathrm{InSM}}$ | half-saturation constant for inhibition of $S$ by $M$ | 0.125 | AU/AV |
| $k_{\mathrm{TfIn}}^{\mathrm{As}}$ | association of transcription factor and inhibitor | 200 | AV/(AU·min) |
| $k_{\mathrm{M}}^{\mathrm{De}}$ | constitutive degradation of $M$ | 0.1 | 1/min |
| $k_{\mathrm{MA}}^{\mathrm{De}}$ | degradation of $M$ by antagonist | 10 | AV/(AU·min) |
| $k_{\mathrm{S}}^{\mathrm{De}}$ | constitutive degradation of $S$ | 0.1 | 1/min |
| $k_{\mathrm{Sk}}^{\mathrm{De}}$ | constitutive starter kinase degradation | 0.0018 | 1/min |
| $k_{\mathrm{SkLi}}^{\mathrm{De}}$ | light-dependent starter kinase degradation | 0.021 | 1/min |
| $k_{\mathrm{A}}^{\mathrm{Dp}}$ | dephosphorylation of antagonist | 2 | 1/min |
| $k_{\mathrm{In}}^{\mathrm{Dp}}$ | dephosphorylation of inhibitor | 2 | 1/min |
| $k_{\mathrm{TfIn}}^{\mathrm{Ds}}$ | dissociation of transcription-factor inhibitor complex | 0.2 | 1/min |
| $k_{\mathrm{AM}}^{\mathrm{Ph}}$ | phosphorylation of antagonist by $M$ | 2000 | $AV^2/(AU^2 \cdot min)$ |
| $k_{\mathrm{AS}}^{\mathrm{Ph}}$ | phosphorylation of antagonist by $S$ | 25 | $AV^2/(AU^2 \cdot min)$ |
| $k_{\mathrm{InFk}}^{\mathrm{Ph}}$ | phosphorylation of inhibitor by feedback kinase | 7.4 | 1/(AU·min) |
| $k_{\mathrm{InSk}}^{\mathrm{Ph}}$ | phosphorylation of inhibitor by starter kinase | 0.05 | AV/(AU·min) |
| $k_{\mathrm{MM}}^{\mathrm{Sy}}$ | synthesis of $M$ promoted by $M$ | 0.06 | 1/min |
| $k_{\mathrm{MS}}^{\mathrm{Sy}}$ | synthesis of $M$ promoted by $S$ | 0.2 | 1/min |
| $k_{\mathrm{Sk}}^{\mathrm{Sy}}$ | starter kinase synthesis | 0.015 | $AU/(AV^2 \cdot min)$ |
| $k_{\mathrm{STf}}^{\mathrm{Sy}}$ | transcription factor-dependent synthesis of $S$ | 0.25 | 1/min |
| $Light$ | binary variable indicating the presence of light | 0 or 1 | – |
| $n_{\mathrm{InSM}}$ | hill exponent for inhibition of $S$ by $M$ | 4 | – |
| $n_{\mathrm{PhA}}$ | hill exponent for phosphorylation of antagonist | 2 | – |
| $TF_{\mathrm{t}}$ | total transcription factor concentration | 1 | AU/AV |

[a]AU, arbitrary unit of number of molecules; AV, arbitrary unit of cell volume.

**Table S2. Non-zero initial conditions for multiple fission model.**

| Variable | Description | Value | Unit[a] |
|---|---|---|---|
| $V$ | cell volume | 1 | AV |
| $SK$ | starter kinase concentration | 8.3 | AU/AV |

[a]AU, arbitrary unit of number of molecules; AV, arbitrary unit of cell volume.

**Table S3. Parameter changes for perturbation simulations in Fig. 4 and S4.**

| Label | Parameter | Reduction (-) | Increase (+) | Unit[a] |
|---|---|---|---|---|
| FK | $k_{\mathrm{InFk}}^{\mathrm{Ph}}$ | 3.7 | 14.8 | 1/(AU·min) |
| IN | $IN_{\mathrm{t}}$ | 1.4 | 2.6 | AU/AV |
| Li | $k_{\mathrm{SkLi}}^{\mathrm{De}}$ | 0.0021 | 0.21 | 1/min |
| SK | $k_{\mathrm{Sk}}^{\mathrm{Sy}}$ | 0.0015 | 0.0285 | AU/(AV$^2$·min) |
| TF | $TF_{\mathrm{t}}$ | 0.7 | 1.3 | AU/AV |

[a]AU, arbitrary unit of number of molecules; AV, arbitrary unit of cell volume.
